# AMP-activated protein kinase is essential for the maintenance of energy levels during synaptic activation

**DOI:** 10.1101/303867

**Authors:** Claudia Marinangeli, Sébastien Didier, Tariq Ahmed, Raphaelle Caillerez, Manon Domise, Charlotte Laloux, Séverine Bégard, Sébastien Carrier, Morvane Colin, Philippe Marchetti, Bart Ghesquière, Detlef Balschun, Luc Buée, Jérôme Kluza, Valérie Vingtdeux

## Abstract

While accounting for 2% of the total body mass, the brain is the organ that consumes the most energy. Although it is widely acknowledged that neuronal energy metabolism is tightly regulated, the mechanism how neurons meet their energy demand to sustain synaptic transmission remains poorly studied. Here we provide substantial evidence that the AMP-activated protein kinase (AMPK) plays a leading role in this process. Our results show that following synaptic activation, AMPK activation is required to sustain neuronal energy levels particularly through mitochondrial respiration. Further, our studies revealed that this metabolic plasticity regulated by AMPK is required for the expression of immediate early genes, synaptic plasticity and memory formation. These findings are important in the context of neurodegenerative disorders, as AMPK deregulation as it is observed in Alzheimer’s disease, impairs the metabolic response to synaptic activation. Altogether, our data provides the proof of concept that AMPK is an essential player in the regulation of neuroenergetic metabolism plasticity induced in response to synaptic activation.

## Introduction

Besides accounting for only 2% of the total body mass, the brain is one of the most energy consuming organs, requiring up to 20 % of the oxygen and 25% of the glucose metabolized by the body. These substrates are converted to ATP through glycolysis and oxidative phosphorylation, the latter taking place in mitochondria. Hence, around 20% of the ATP produced in the body is devoted to the brain. Importantly, brain functions depend on an adequate energy supply. Indeed, reducing the brain supply of glucose or oxygen even for short periods leads to severe brain damages as observed following ischemic stroke. Numerous reports support a role for glucose as a potent modulator of learning and memory including behavioral tasks in both humans and rodents. Indeed, both peripheral and direct central glucose administration enhances cognitive processes (Gold, 1995). Similarly, astrocytes derived lactate, the end product of glycolysis, was reported to be necessary for the formation of long-term memory (Suzuki et al, 2011). Consistent with these observations, brain imaging approaches are based on the detection of signals that are directly related to energy supply and use. For instance, positron emission tomography (PET) reveals changes in cerebral blood flow, glucose uptake and oxygen consumption according to the tracer that is used. These imaging approaches have been instrumental in demonstrating that brain activity is coupled to increased energy availability and consumption. Further studies have demonstrated that 85 % of the energy required by the brain is used by neurons and is devoted to glutamatergic transmission (Attwell & Laughlin, 2001; Harris et al, 2012).

Failure to maintain proper energy levels in the brain might also lead to neurodegeneration. The main neurodegenerative diseases are characterized by deregulations of the energy metabolism (Ferreira et al, 2010). For instance, Alzheimer’s disease (AD) is characterized by mitochondrial dysfunctions (Lin & Beal, 2006) and early glucose hypometabolism (Mosconi, 2005). Of note, [^18^F] fluorodeoxyglucose (FDG)-PET, a marker for glucose utilization, was recently described as being the most accurate biomarker for AD. Indeed, FDG-PET allowed to correctly differentiate subjects who converted to AD or other dementia from those who remained stable or reverted to normal cognition (Caminiti et al, 2018). Finally, while these metabolic impairments were proposed to play a causative role in the disease pathogenesis, their causes and consequences remain to be established.

One of the most important cell energy sensors and regulators is the AMP-activated protein kinase (AMPK). AMPK is a serine/threonine protein kinase composed of three subunits. It possesses one catalytic subunit alpha and two regulatory subunits beta and gamma. The gamma subunit holds adenine nucleotide binding sites that allow the sensing of intracellular levels of AMP, ADP and ATP. Consequently, any energetic stress that will modify the intracellular AMP/ATP ratio will thereby enhance the activation of AMPK (Xiao et al, 2007). The activation of AMPK requires its phosphorylation on threonine 172 within its catalytic alpha subunit. Two main kinases were reported to fulfill this role, the liver kinase B1 (LKB1), which is thought to be constitutively activated and involved in AMPK phosphorylation in energy stress conditions (Hawley et al, 2003; Woods et al, 2003), and the Ca^2+^/calmodulin-dependent protein kinase kinase-β (CaMKKβ), whose activation is dependent on intracellular calcium levels (Hawley et al, 2005). AMPK is highly expressed in the brain and in particular in neurons. Interestingly, several studies have reported an over-activation of neuronal AMPK in the brain of Alzheimer’s (Vingtdeux et al, 2011), Parkinson’s (Jiang et al, 2013), and Huntington’s (Ju et al, 2011) patients, thus supporting the defective metabolism hypothesis in these disorders (Domise & Vingtdeux, 2016).

While it is obvious that brain energy metabolism must be tightly regulated to sustain neuronal function and cognition, the molecular mechanisms involved in this regulation remain poorly understood. In particular, how neurons meet their energy demand to sustain synaptic transmission remains to be established. In this study, we assessed the possibility that AMPK might play a critical role in the regulation of energy metabolism in neurons in response to their activation.

## Results

### AMPK is stimulated following synaptic activation

To assess the molecular pathways involved in neuronal energy levels maintenance, we used bicuculline and 4-aminopyridine (Bic/4-AP) to induce synaptic release of glutamate in differentiated primary neurons. This well-established protocol (Hardingham et al, 2002; Hoey et al, 2009) is known to induce preferentially glutamatergic neurotransmission, that we will refer hereafter as synaptic activation (SA). To validate the efficiency of this protocol in our model, we first assessed the activation status of a signaling pathway known to be activated following SA, the MAPK pathway. SA induction in 15 *days in vitro* (DIV) differentiated mouse primary neurons led to a rapid phosphorylation of the main MAPK pathway components, ERK, RSK, MSK and MEK, which are representative of this signaling pathway activation (Fig. EV1 A). In addition, blockade of glutamatergic NMDA and AMPA repectors by the specific inhibitors MK-801 and NBQX respectively, completely abolished the activation of ERK, confirming that MAPK activation is dependent on the stimulation of glutamate receptors (Fig. EV1 B). Together, these results confirm the validity of the SA protocol in our model. To allow a more general screening of the signaling pathways activated following SA we used a kinase array approach. After 5 min Bic/4-AP stimulation, we detected an increased in the phosphorylation status of ERK, MSK, and CREB as it was expected from our previous data (Fig. 1 A-B, E). Interestingly, an increase in the phosphorylation levels of AMPK was also observed following Bic/4-AP treatment. AMPK activation was further assessed by Western-blotting (WB) using phospho-specific antibodies directed against the AMPK-Thr^172^ epitope, that is a prerequisite for AMPK activity, and Acetyl-CoA carboxylase (ACC)-Ser^79^, a direct AMPK target (Fig. 1 G-I). Following SA, we observed a rapid phosphorylation of AMPK and ACC thus demonstrating that the AMPK signaling pathway was activated (Fig. 1 G-I). This activation was also dependent on glutamate receptors activation since their inhibition significantly reduced AMPK activation (Fig. EV1 B-D). Given that SA leads to intracellular calcium influx and that AMPK was reported to be phosphorylated by the calcium-dependent protein kinase CaMKKβ in neuronal cells, we used the CaMKKβ specific inhibitor STO-609 to address this possibility. However, our results showed that in our model, CaMKKβ was not responsible for the SA-induced AMPK activation (Fig. EV2) thus strongly suggesting that AMPK phosphorylation was not related to intracellular calcium levels. Since SA is a very energetic process, it is likely that AMPK activation might be related to changes in ATP levels. To assess the importance of energy levels maintenance for the signaling pathways induced by SA, primary neurons were pretreated with a combination of oligomycin, an ATP synthase inhibitor, and 2-deoxy-D-glucose, a glycolysis inhibitor (Oligo/2DG) in order to deplete intracellular ATP (Fig. 1 F). In these conditions of energy depletion, SA failed to activate the MAPK signaling pathway (Fig. 1 C-E). However, AMPK activity was increased in both conditions (Fig. 1 C-E). Altogether, our data show that the activation of the MAPK signaling pathway by SA is contingent upon intracellular energy levels suggesting a role of AMPK in their regulation.

**Figure 1.**
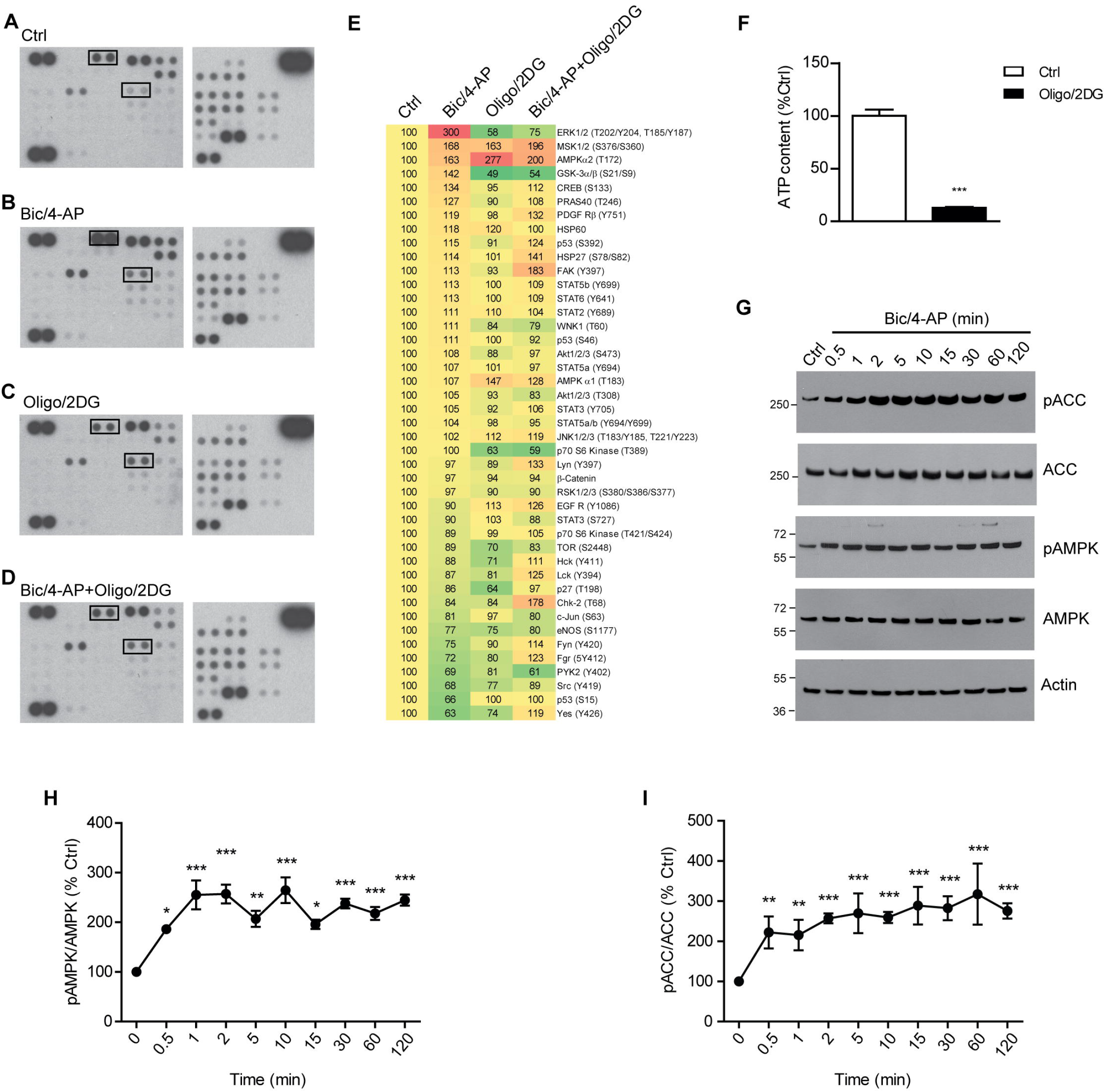
AMPK is activated upon synaptic activation. A-D Total lysates from 15 DIV *(days in vitro)* differentiated primary neurons stimulated with Bic/4-AP (5 min) in presence or absence of oligomycin and 2-deoxy-D-glucose (Oligo/2DG, 5 min pretreatment 1 μM/50 mM) were subjected to a kinase array (A to D, upper boxes show phosphorylated ERK1/2 (Thr^202^/Tyr^204^, Thr^185^/Tyr^187^) and lower boxes show phosphorylated AMPK (Thr^172^)). E Quantification of the kinase array represented as a heat map, results are expressed as percentage of the Ctrl. F Intracellular ATP quantifications following 5 min Oligo/2DG treatment (n=3). G Western-blotting (WB) analysis of phosphorylated ACC and AMPK and total ACC, AMPK, and Actin in lysates obtained from 15 DIV neurons stimulated with Bic/4-AP for the indicated times. WB are representative of at least 4 independent experiments. H-l WB quantifications showing the ratios of phosphorylated AMPK/total AMPK (pAMPK/AMPK) (H) and phosphorylated ACC/total ACC (pACC/ACC) (I) following Bic/4-AP stimulation (n=4). Results show means ± SD. Student’s t test (F) and One-way ANOVA with Bonferonni post hoc (H-l) were used for evaluation of statistical significance. * p < 0.05, ** p < 0.01, *** p < 0.001.

### AMPK activation is necessary to induce neuronal metabolic plasticity in response to synaptic activation

To test whether AMPK was involved in the maintenance of energy levels during SA, we assessed metabolic fluxes in differentiated live neurons using the extracellular flux analyzer Seahorse XFe24 (Marinangeli et al, 2018). Glycolytic flux was determined by measuring the extracellular acidification rate (ECAR) due to H^+^ release in combination with lactate, and mitochondrial respiration was determined by measuring the oxygen consumption rate (OCR). Subsequent to Bic/4-AP stimulation, ECAR and OCR were increased showing that both glycolysis and mitochondrial respiration were enhanced to maintain energy levels following SA (Fig. 2 A and G, green arrows, and Fig. 2 C, I). Several metabolic parameters were determined using calculations described in the methods section. Interestingly, Bic/4-AP stimulation did not change the overall glycolytic reserve (Fig. 2 E), maximal respiration (Fig. 2 K), or spare respiratory capacity (Fig. 2 L) but rather took on the existing glycolytic reserve (Fig. 2 F) and spare respiratory capacity (Fig. 2 M) to increase energy production (Fig. 2 J). To determine the involvement of AMPK in this metabolic up-regulation induced following SA, neurons were pretreated with the AMPK inhibitor Compound C (Cc). Our results showed that AMPK inhibition prohibited the up-regulation of glycolysis and mitochondrial respiration that were induced by SA (Fig. 2 A-M). While AMPK inhibition in basal conditions (without SA induction) did not affect overall basal glycolysis or mitochondrial functions (Fig. EV3), it significantly reduced the use of the spare glycolytic reserve and spare respiratory capacity induced by SA (Fig. 2 F and M). Similar results were obtained using a kinase dead dominant negative construct of AMPK therefore excluding any potential off-target effect of Cc (Fig. EV4 A-B). Finally, the phenogram analysis, that defines the overall cellular metabolic state of the cells, demonstrated that synaptic activation induced by Bic/4-AP led to a shift towards an energetic state of the neurons (Fig. 2 N) that we will refer to as their metabolic plasticity. AMPK inhibition using Cc almost completely abolished this shift (Fig. 2 N), thereby suggesting that AMPK activity is necessary to maintain energy levels during SA. To confirm this, intracellular ATP levels were quantified in neurons at different time points following SA in presence or absence of Cc. While in control and SA conditions ATP levels remained constant, AMPK repression led to a significant rapid drop in ATP levels in conditions of SA (Fig. 2 O). These results showed that AMPK activity following SA is essential to maintain neuronal energy levels.

**Figure 2.**
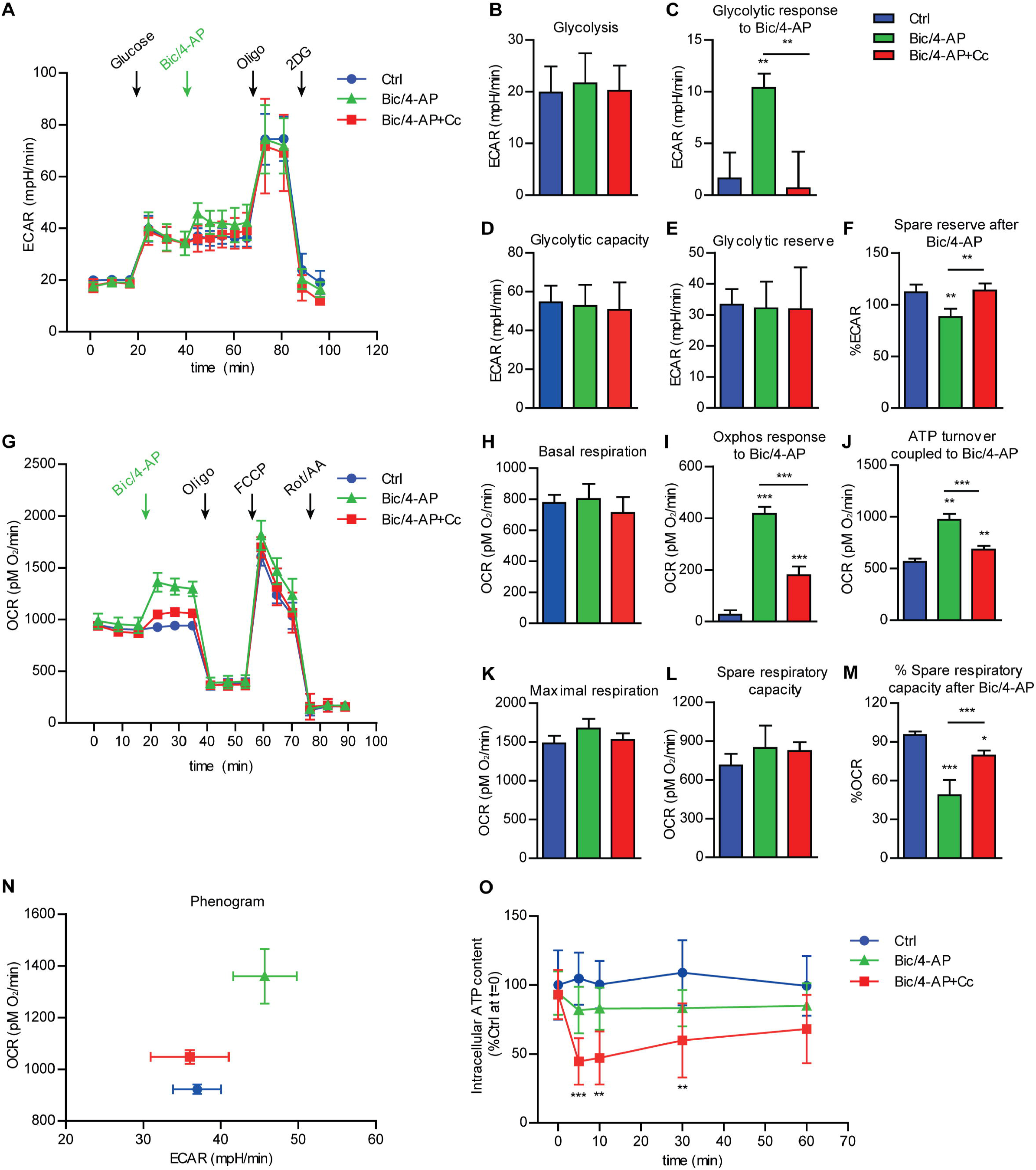
AMPK maintains energy levels by up-regulating glycolysis and mitochondrial respiration during synaptic activity. A-M (A) Extracellular acidification rate (ECAR) and (G) oxygen consumption rate (OCR) measured using the Seahorse technology after Bic/4-AP stimulation (green arrows) in the presence or absence of the AMPK inhibitor Compound C (Cc, 10 μM). (A) ECAR profile is monitored under basal conditions and upon the sequential injection of saturating concentration of glucose, followed by Bic/4-AP, or assay medium in Ctrl group, oligomycin (Oligo) and 2-deoxy-D-glucose (2DG) as indicated by arrows. (G) OCR profile monitored under basal condition and following the sequential injection of assay medium, in Ctrl group, or Bic/4-AP, in the stimulated groups, Oligo, FCCP, rotenone and antimycin A (Rot/AA) as indicated by arrows. ECAR and OCR are indicators of glycolysis and mitochondrial respiration, respectively. (B) Basal glycolysis, (C) glycolytic response to Bic/4-AP, (D) glycolytic capacity, and (E) spare glycolytic reserve expressed as mpH/min, (F) spare glycolytic reserve after Bic/4AP expressed as percentage of baseline, (H) basal respiration, (I) oxphos response to Bic/4AP, (J) ATP turnover coupled to Bic/4AP stimulation, (K) maximal respiration, (L) spare respiratory capacity expressed as pMO_2_/min, and (M) spare respiratory capacity after Bic/4-AP stimulation expressed as percentage of baseline were calculated as described in the methods section n=4-5. N Phenogram obtained by plotting the values obtained from the study of glycolysis (A) and mitochondrial respiration (G) following Bic/4-AP stimulation. O Intracellular ATP levels in neurons stimulated with Bic/4-AP for the indicated times in presence or absence of Cc (n=3-4). Values are expressed as percentage of ATP in the Ctrl at time O. Results show mean ± SD. One-way ANOVA (B-F and H-M) and two-way ANOVA (O) followed by Bonferroni’s post-test were used for evaluation of statistical significance. * p < 0.05, ** p < 0.01, *** p < 0.001.

### AMPK up-regulates mitochondrial respiration in response to synaptic activation

Glycolysis and mitochondrial respiration are tightly coupled together. Given that neurons are mainly oxidative cells, we asked whether the increase in mitochondrial respiration induced by SA was dependent on glycolysis up-regulation or whether AMPK could have a more direct impact on mitochondria. To assess the latter, we investigated its involvement in the oxidation of mitochondrial substrates that do not require prior glycolysis. As neurons were reported to use lactate as an alternative substrate to glucose in the context of the astrocyte-neuron lactate shuttle, we substituted glucose for lactate in the Seahorse experiments. In these conditions, SA up-regulated mitochondrial respiration (Fig. EV5 A-F) as it was previously observed with glucose. This metabolic response to SA was inhibited by the presence of Cc, showing that AMPK is also involved in the regulation of lactate utilization following SA. Importantly, these results support a more direct role of AMPK in mitochondrial function. Given that neurons were also reported to use glutamine for energetic purpose (Divakaruni et al, 2017), we repeated these experiments with a media devoid of glucose. Results showed that following SA, neurons did not use the glutamine present in the culture media to increase their energy production (Fig. EV5 G-L). To go further, we performed metabolomics to determine fluxes throughout central carbon metabolism using ^13^C stable-isotope tracing. For this, neurons were cultured in the presence of uniformly ^13^C-labeled glucose and fractional contribution was quantified using mass spectrometry (Isotopologues were designated as mO, ml, m2, m3…). An increase in labeled lactate confirmed the increased glycolytic flow under SA conditions that was prohibited by Cc (Fig. 3 A). Furthermore, analysis of the isotopologue profiles of the tricarboxylic acid cycle (TCA) metabolites demonstrated significant changes (Fig. 3 B-H). Of particular interest is the significant increase in the levels of m3 isotopomers in SA from citrate, succinate, fumarate and malate, suggesting that SA induced a shift towards pyruvate carboxylation (Buescher et al, 2015). Higher abundances of m4, m5 and m6 isotopologues from citrate were observed following Bic/4-AP stimulation (Fig. 3 C). These isotopomers require the combined action of pyruvate dehydrogenase (PDH), pyruvate carboxylase (PC), and multiple turns of the Kerbs cycle as explained in Fig. 3B. These results suggest that Bic/4-AP stimulation induced a shift towards anaplerotic pyruvate carboxylation that was not observed when AMPK activity was repressed. Altogether, these data support the notion that AMPK activity is needed to induce the metabolic “reprogramming” that occurs following SA.

**Figure 3.**
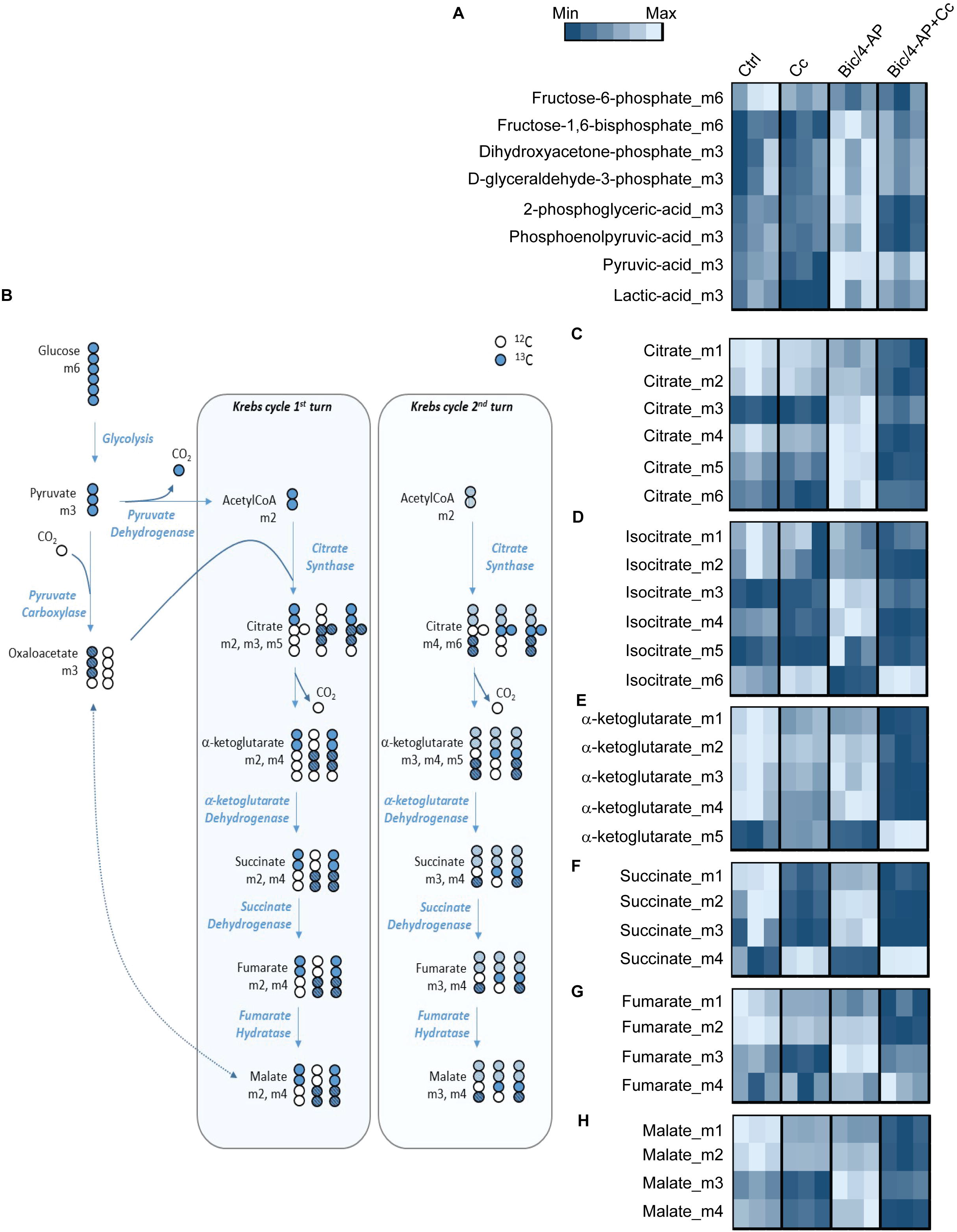
AMPK up-regulates mitochondrial metabolism upon synaptic activation. Measurement of ^13^C-glucose derived metabolites, calculated as percentage of the total metabolite pool following LC-MS analysis of primary neurons after 30 min in culture with ^13^C-glucose-supplemented media with or without Bic/4-AP stimulation. A Metabolic profile of glycolysis intermediates. B Schematic representation of the labeling patterns of metabolic intermediates starting from uniformly labeled glucose. The letter “m” indicates the number of carbon atoms labeled with ^13^C. The blue filled circles represent ^13^C atoms derived from pyruvate dehydrogenase activity while the dashed circles represent ^13^C atoms derived from the pyruvate carboxylase activity. C-H Isotopologues quantification of the Krebs cycle intermediates citrate (C), isocitrate (D), α-ketoglutarate (E), succinate (F), fumarate (G) and malate (H). Metabolic profiles are expressed as heat maps. n=3.

### Neuronal metabolic plasticity induced by AMPK is crucial for immediate early genes expression, synaptic plasticity and long-term memory formation

We next sought to determine the importance of this metabolic plasticity regulated by AMPK for neuronal functions. Synaptic activation, when sustained, leads to the expression of immediate early genes (IEGs) that include Arc, cFos and Egr1. These genes are known to modulate the neuronal plasticity underlying learning and memory. In our model of synaptic activation, we observed the expression of these IEGs starting 1 hr after Bic/4-AP treatment (Fig. 4 A). This induction of IEGs expression was inhibited when the energetic status was disrupted by either inhibition of glycolysis with the glyceraldehyde-3-phosphate inhibitor iodoacetate or 2-DG or by interruption of mitochondrial respiration using oligomycin or the complexes I and III inhibitors rotenone and antimycin A, respectively (Fig. 4 B). These results demonstrated that the maintenance of energy levels is crucial for the expression of IEGs. Additionally, AMPK inhibition also led to a significant reduction of IEGs expression following SA (Fig. 4 C and EV4 C), showing that AMPK-mediated metabolic plasticity is necessary for the expression of IEGs following SA.

**Figure 4.**
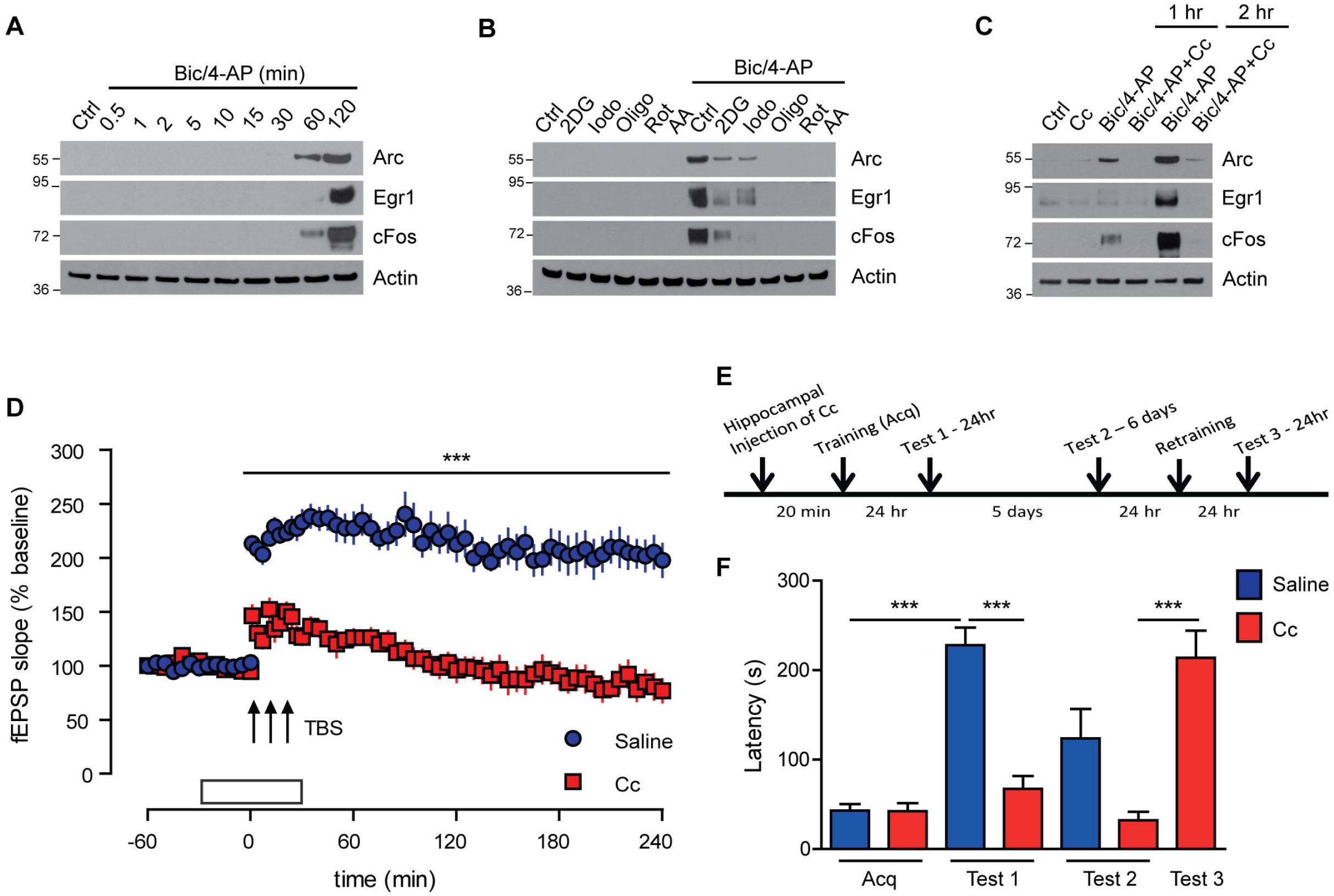
AMPK activation following synaptic activity is necessary for IEGs expression, synaptic plasticity and long-term memory formation. A WB analysis of the IEGs Arc, Egrl and cFos expression following Bic/4-AP stimulation for the indicated times in differentiated primary neurons at 15 DIV. WB are representative of at least 5 independent experiments. B WB analysis of the IEGs Arc, Egrl, and cFos expression following 2 hr Bic/4-AP stimulation in the presence or absence of glycolysis inhibitors (2DG 50 mM, lodo 100 μM pre-treatment of 5 min) or mitochondrial respiration inhibitors (Oligo 1 μM, Rot 1 μM, AA lμM pre-treatment of 5 min) in differentiated primary neurons at 15 DIV. WB are representative of at least 4 independent experiments. C WB analysis of the IEGs Arc, Egrl and cFos expression following 2 hr Bic/4AP stimulation in the presence or absence of Cc (20 min pretreatment, 10 μM). WB are representative of at least 4 independent experiments. D Recording of LTP in the CAl-region of the hippocampus *ex vivo.* Triple application of a theta-burst electrical stimulation protocol (TBS) induced robust LTP in controls. However, in slices treated with lOμM Cc, the same stimulation generated only an impaired, decremental potentiation, returning to baseline values after about 80 min (Wilcoxon-test). This resulted in a highly significant difference (Fı ı_3_=120.105; p<0.001, RM-ANOVA). Data are presented as mean ± SEM, where n refers to the number of animals tested. E Schematic representation of the Inhibitory avoidance (IA) protocol timeline. F Hippocampal injections of Cc 20 min before the IA training (Acq) disrupted long-term memory at 24 h (Test 1). The disruption persisted 6 days after training (Test 2). Cc injected mice had normal retention after retraining (Test 3). n=10, results show mean ± SEM. One-way ANOVA followed by Bonferroni’s post-test was used for evaluation of statistical significance. * p < 0.05, ** p < 0.01, *** p < 0.001.

In order to examine the impact of AMPK inhibition on synaptic plasticity under more physiological conditions, we took advantage of *ex vivo* hippocampal slices. Thus, we investigated a putative role of AMPK in long-term potentiation (LTP) in the CA1-region of the hippocampus, a well-established model of learning and memory at the cellular level. As depicted in Fig. 4D, triple application of a theta-burst electrical stimulation protocol (TBS) resulted in robust LTP in controls with initial values of 213.3 ± 5.3 % (n=7) which were retained at about this level until 4 hours after induction (197.5 ± 15.1 %). This robust type of LTP is protein-synthesis dependent and therefore a relevant model for the formation of long-term memory (Ahmed et al. submitted). Application of 10 μM Cc, in contrast, caused a severe impairment of LTP with a significantly lower initial magnitude of 146.2 ± 9.4 % (p=0.0001 Welch-test) and a decremental potentiation that returned to baseline values after about 80 min (Wilcoxon-test). Overall, this resulted in a highly significant difference to LTP in control mice (F_1,13_=120.105; p<0.001, RM-ANOVA) underlining an essential function of AMPK in long-term synaptic plasticity.

Finally, to evaluate the importance of AMPK-mediated metabolic plasticity *in vivo,* we tested whether AMPK activation is involved in long-term memory retention. To this end, Cc was bilaterally injected in the hippocampus of awake wild-type mice 20 min before inhibitory avoidance training (IA), a task that depends on the proper functioning of the hippocampus (Izquierdo et al, 2016; Whitlock et al, 2006) (Fig. 4 E-F). Cc injection did not affect IA acquisition, as the mean latencies to enter the shock compartment during training (Acq) were similar in both groups. However, Cc significantly blocked long-term memory tested at 24 hr (Test 1). Retesting 6 days after acquisition (Test 2) showed that the memory loss persisted. Retraining of the Cc injected group, resulted in normal memory retention 24 hr later (Test 3), indicating that the hippocampus was functionally intact. Altogether, these results show that AMPK is involved in long-term memory formation and provided the proof-of-concept that AMPK plays an essential role in cognitive processes.

### AMPK deregulation impairs the neuronal metabolic plasticity induced by synaptic activation

Lastly, we asked whether AMPK deregulation as it is observed in AD could impair AMPK physiological function. Since AMPK is found overactivated in neurons of AD patients as compared to control patients (Vingtdeux et al, 2011), we mimicked this condition and activated AMPK by taking advantage of AICAR, an AMPK agonist which was applied 24 hr before SA stimulation (referred to as preactivation hereafter). While, AMPK pre-activation with AICAR did not modify the metabolic profile of neurons, AMPK pre-activation greatly impaired the ability of neurons to up-regulate mitochondrial respiration in response to Bic/4-AP (Fig. 5 A-B). Similar results were obtained after over-expression of a constitutively active mutant of AMPK (CA-AMPK) in neurons (Fig. 5 C) further supporting the inhibitory effect of AMPK pre-activation on the metabolic response to SA. Finally, our results show that the pre-activation of AMPK either using AICAR or CA-AMPK prevented the SA-induced expression of IEGs (Fig. 5 D-E). Altogether, our results suggest that a tight regulation of AMPK is important for the physiological metabolic response to SA to occur and for the triggering of processes leading to neuronal plasticity.

**Figure 5.**
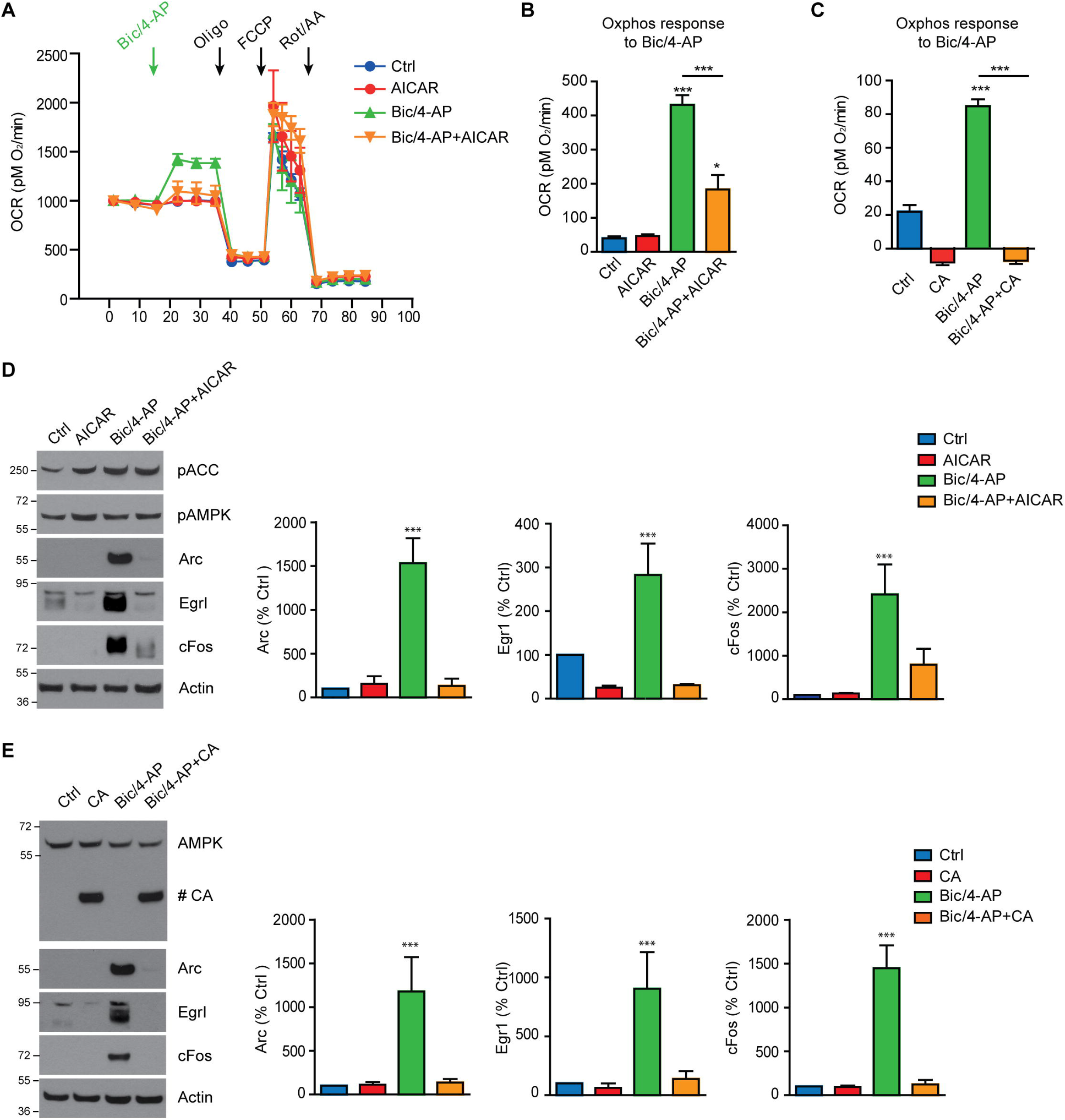
AMPK pre-activation prevents neuronal metabolic response to SA. A Mitochondrial respiration profile in primary neurons evaluated using the Seahorse technology to investigate the impact of long lasting (24 hr) AMPK activation induced by AICAR in response to Bic/4-AP. Mitochondrial respiration profile, expressed as oxygen consumption rate (OCR), is monitored upon the injection of assay medium, in control group (Ctrl) and in cells treated with AICAR (AICAR), or upon the injection of Bic/4-AP in absence (Bic/4-AP) or presence of AICAR (Bic/4-AP+AICAR), followed by injection of Oligo, FCCP and Rot/AA as indicated by the arrows. Results are representative of 3 independent experiments. B OCR response to Bic/4-AP stimulation expressed as pMO_2_/min (n=3). C OCR response to Bic/4-AP stimulation expressed as pMO_2_/min in primary neurons overexpressing a constitutively active (CA) form of AMPK. (n=3). D Immunoblotting with phosphorylated ACC (pACC), phosphorylated AMPK (pAMPK), Arc, Egrl, cFos, and actin in lysates obtained from 15 DIV neurons stimulated with Bic/4-AP for 2 hr after pre-treatement with AICAR (24h, 1mM). Results are representative of at least 5 independent experiments. Quantification of WB showing the expression of the IEGs Arc, Egrl and cFos (n=5). E Immunoblotting with total AMPK (# CA labels the overexpressed CA-AMPK construct), Arc, Egrl, cFos, and Actin in lysates obtained from 15 DIV neurons stimulated with Bic/4-AP for 2 hr, 4 days after infection with CA-AMPK. Results are representative of at least 3 experiments. Quantification of WB showing the expression of the IEGs Arc, Egrl and cFos (n=3). Results show mean ± SD. One-way ANOVA followed by Bonferroni’s post-test was used for evaluation of statistical significance. * p < 0.05, ** p < 0.01, *** p < 0.001).

## Discussion

Neurons have high energy demands that are met through the oxidation of glucose via glycolysis and mitochondrial respiration. Neuronal activity is a process known to require and thus stimulate ATP production (Rangaraju et al, 2014), however the molecular mechanisms involved in this regulation remain poorly understood. Here, we demonstrated that the AMPK-mediated signaling pathway plays a key role in this regulation. We found that AMPK is rapidly activated following synaptic activation to maintain energy levels via the up-regulation of glycolytic and mitochondrial use of glucose. Consistent with their high energy requirements, neurons sustain a high rate of oxidative metabolism (Boumezbeur et al, 2010). Bearing this in mind, the importance of mitochondria for cognitive functions has recently regained importance. In particular with a recent study showing that cannabinoid receptors present on mitochondria (mtCB1) are responsible for amnesia induced by acute cannabinoid intoxication. In particular, activation of hippocampal mtCB1 receptors led to memory impairments due to decreased mitochondrial respiration (Hebert-Chatelain et al, 2016). Our study reinforce this link between mitochondrial activity and memory formation. Our results also raised the question of AMPK subcellular localization. Indeed, it could be speculated that a pool of AMPK might be present at or within the mitochondria to up-regulates mitochondrial respiration. Previous reports have also highlighted this possibility (Liang et al, 2015). In our context, further work will be required to determine whether PC or PDH could be direct AMPK substrates. However, another interesting possibility is the regulation by AMPK of the AKAP1 (A kinase anchor protein 1) protein, that was identified in muscle by a global phosphoproteomic approach (Hoffman et al, 2015). This direct phosphorylation of AKAP1 by AMPK was reported to regulate mitochondrial respiration (Hoffman et al, 2015). While we assessed only the immediate metabolic response of neurons to synaptic activity, it will also be interesting to determine whether AMPK could be involved in gene expression changes that promotes the neuronal Warburg effect in response to prolonged Bic/4-AP stimulation (Bas-Orth et al, 2017).

Our data also demonstrate that the energetic metabolic up-regulation mediated by AMPK is crucial for the expression of IEGs, synaptic plasticity and hence memory formation. In line with these results, AMPK was reported to be activated and necessary for LTP induced *in vivo* by high-frequency stimulation (Yu et al, 2016). However, for a more thorough understanding of these processes, the generation of conditional AMPK knock-out mouse models would help in elucidating the importance of AMPK in cognitive processes.

Finally, our results are also particularly important in the context of neurodegenerative disorders in which energy metabolism is strongly impaired and AMPK overactivated (Jiang et al, 2013; Ju et al, 2011; Vingtdeux et al, 2011). In conditions of overactivation, AMPK is unlikely to respond properly to synaptic activation, perhaps as a way to preserve energy levels. As a consequence, we propose that AMPK might act as a metabolic switch that determines whether synaptic activation may or may not occur according to the currently available energy levels (and hence metabolic substrate) in its vicinity. It is also possible that AMPK over-activation could indirectly impair the metabolic plasticity response to synaptic activation. Under these conditions, the identification of AMPK targets as well as the consequences of AMPK over-activation on synaptic integrity will be of interest. Supporting this hypothesis, previous reports have determined that AMPK activation by pharmacological drugs led to impairments in synaptic plasticity (Potter et al, 2010) and memory formation (Dash et al, 2006). In addition, in the context of AD, AMPK was proposed to be a key player in the reduction of dendritic spines numbers (Mairet-Coello et al, 2013) and in the LTP impairments induced by Aβ oligomers (Ma et al, 2014). These reports show that upregulation of AMPK activity has the same negative outcome on synaptic plasticity and memory formation as AMPK inhibition. Altogether, our data suggest that AMPK activity has to be extremely well fine-tuned since any hypo- or hyper-activation will have detrimental consequences on synaptic plasticity and hence memory formation.

In summary, our results demonstrate that AMPK plays a critical role during glutamatergic synaptic activation. In particular, we show that AMPK activity is crucial to maintain neuronal energy levels and hence is involved in memory formation processes. Our study establishes AMPK as an important link between memory formation and energy metabolism and suggests that it could act as a metabolic checkpoint by sensing energy substrates availability. These findings might also be of value in the context of diabetes and obesity, conditions that are associated with cognitive (Dye et al, 2017; McCrimmon et al, 2012) and central metabolism (Hwang et al, 2017) impairments in humans.

## Material and methods

### Chemicals and reagents/antibodies

Oligomycin, Carbonyl cyanide-4-(trifluoromethoxy)phenylhydrazone (FCCP), Antimycin A (AA), Rotenone (Rot), 4-Amynopyridine (4-AP), L-Lactate (Lac), iodoacetate and sodium pyruvate (NaPyr) were purchased from Sigma, 2-deoxy-D-glucose (2DG) from Acros, Compound C (Cc) from Santa Cruz, Bicuculline (Bic), MK-801, NBQX, STO-609, N1-(β-D-Ribofuranosyl)-5-aminoimidazole-4-carboxamide (AICAR) and Metformine from Tocris, Glucose 20% solution was purchased from Invitrogen. Antibodies directed against AMPKα, ACC, phospho-Ser^79^ACC, ERK1/2, phospho-Thr^202^/Tyr^204^ERKl/2, phospho-Thr^581^MSK, phospho-Thr^380^p90RSK and phospho-Ser^133^CREB were obtained from Cell Signaling technology. Anti phospho-Thr^172^AMPKα, Arc, cFos, and Egrl antibodies were from Santa-cruz. Anti-actin antibody was from BD Transduction Laboratory.

### Animals

All animal experiments were performed according to procedures approved by the local Animal Ethical Committee following European standards for the care and use of laboratory animals (agreement APAFIS#4689-2016032315498524 v5 from CEEA75, Lille, France).

### Surgical procedures and injections

Three-month-old male C57BL/6J mice were obtained from the Jackson Laboratory. Bilateral hippocampal surgeries were performed as described by Brouillette et al (Brouillette et al, 2012). Briefly, bilateral cannulae (3280PD-2.8/Spc with a removable dummy wire; Plastics One) were stereotaxically implanted into the hippocampus (coordinates with respect to bregma: -2.2 mm anteroposterior [AP], +/-1.4 mm mediolateral [ML], -2.1 mm dorsoventral [DV], according to the Paxinos and Franklin mouse brain atlas) (Paxinos and Watson, 2005) in anesthetized mice (100 mg/kg of ketamine and 10 mg/kg of xylazine, i.p.). After surgery, the animals recovered for 10 days before undergoing any procedures. After 10 days of recovery, awake and freely moving mice were injected with 2 μl of Compound C or an equal volume of vehicle buffer at a rate of 0.25 μl/min via cannulae PE50 tubing (Plastics One) connected to a 10 μL Hamilton syringe pump system (KDS310; KD Scientific). The cannulae was capped to prevent reflux of the injected solution.

### Inhibitory avoidance

Ten days after the surgery, mice underwent the inhibitory avoidance (IA) paradigm. Control mice (Saline) were injected with saline solution (0.9 % NaCl) while the treated mice were injected with Compound C (2 μl at 100 μM in saline) 20 min before running the acquisition test (Acq). The IA apparatus consisted in a rectangular shaped box that is divided into a safe illuminated compartment and a dark shock compartment. During the Acq session, each mouse was placed in the safe compartment and allowed to access the dark chamber where a brief foot shock (0.3 mA, 2 s) was delivered. The mouse was left in the dark compartment for one minute, before being replaced in its home cage. Latency to enter the shock compartment was taken as a measure of acquisition (Acq). 24 hr later, a retention test (Test 1) was run. In this phase, each mouse was placed in the light chamber and the latency to enter the dark compartment was recorded, however no foot shock was delivered when the mouse entered the dark chamber. A second test was run 6 days after the acquisition test (Test 2) in order to assess memory retention. To verify the ability of the injected mice to make new long term memory, each injected mouse underwent a second acquisition test 7 days after the first one (Retraining), and retention was tested 24 hr after (Test 3).

### Extracellular long-term recordings in the CAl-region of the hippocampus

Mice were killed by cervical dislocation and hippocampal slices prepared from the dorsal area of the right hippocampus as reported previously (Ahmed et al, 2015; Denayer et al, 2008). In brief, the right hippocampus was rapidly dissected out into cold (4°C) artificial cerebrospinal fluid (ACSF), saturated with carbogen (95% O_2_/5% CO_2_). ACSF consisted of 124 NaCI, 4.9 KCI, 25.6 NaCO_3_, 1.20 KH_2_PO_4_, 2.0 CaCl_2_, 2.0 MgSO_4_, and 10.0 glucose (in mM), adjusted to pH 7.4. Transverse slices (400 μm thick) were prepared from the dorsal area of the right hippocampus with a tissue chopper and placed into a submerged-type chamber, where they were kept at 32°C and continuously perfused with ACSF at a flow-rate of 2.2 ml/min. After 90 min incubation, one slice was arbitrarily selected and a custom-made tungsten electrode was placed in CA1 stratum radiatum for stimulation in constant current mode. For recording of field excitatory postsynaptic potentials (fEPSPs), a glass electrode (filled with ACSF, 3-7 MΩ resistance) was placed in the stratum radiatum. The time course of the field EPSP (fEPSP) was measured as the descending slope function for all sets of experiments. After a further hour of incubation, input/output curves were established and the stimulation strength was adjusted to elicit a fEPSP-slope of 35% of the maximum and was kept constant throughout the experiment. During baseline recording, three single stimuli (0.1 ms pulse width; 10 s interval) were measured every 5 min and averaged. A robust LTP was induced by three theta burst stimuli (TBS), separated by 10 min. Compound c (Cc) was applied from 30 minutes prior to until 30 min after the first TBS. To allow for direct comparisons all experiments were interleaved between the experimental and control group. Intergroup differences in LTP were examined using ANOVA with repeated measures (RM-ANOVA, SPSS 19). For intragroup comparisons, Wilcoxon’s matched-pairs signed-rank test was employed (SPSS 19). Group differences at single time points were tested using the Welch-test.

### Primary Neuronal Culture

Primary neurons were prepared as previously described (Domise et al, 2016). Briefly, foetuses at stage E18.5 were obtained from pregnant C57BL/6J wild-type female mice (The Jackson Laboratory). Forebrains were dissected in ice-cold dissection medium composed of Hanks’ balanced salt solution (HBSS) (Invitrogen) supplemented with 0.5 % w/v D-glucose (Sigma) and 25 mM Hepes (Invitrogen). Neurons were dissociated and isolated in ice-cold dissection medium containing 0.01 % w/v papain (Sigma), 0.1 % w/v dispase (Sigma), and 0.01 % w/v DNaseI (Roche) and by incubation at 37 °C for 15 min. Cells were spun down at 220 x g for 5 min at 4°C, resuspended in Neurobasal medium supplemented with 2% B27, 1 mM NaPyr, 100 units/ml penicillin, 100 μg/ml streptomycin, 2 mM Glutamax (Invitrogen), and plated at a density of 6*10^6^ cells/plate. Fresh medium was added every 3 days (1:3 of starting volume). Cells were then treated and collected between DIV 14-17.

### Construct and production of lentiviral vectors

N-terminally myc-tagged wild-type (wt) and K45R mutated AMPKα2 rat sequences were subcloned from Addgene plasmids 15991 and 15992 (Mu et al, 2001) into pENTR/D backbone using the In-Fusion HD cloning kit (Clontech) with the following primers: Fwd: 5’-CACCATGGTGCGGGGTTCTCAT 5’-CACCATGGTGCGGGGTTCTCAT and Rev-Trunc: 5’-GTCACCCTAGTATAAACTGTTCATCAC for the wtAMPKα2 and Rev: 5’-TCAACGGGCTAAAGCAGTGATA for the K45R-AMPKα2 to obtain respectively a constitutive active (CA) truncated form of AMPK ending at position 312, the AMPKα2(1-312)-pENTR/D, and a kinase dead dominant negative (DN) form of AMPK, the AMPKα2K45R-pENTR/D. Subsequently, pENTR/D vectors were recombined with SIN-PGK-cPPT-RFA-WHV lentiviral vectors using the Gateway LR Clonase II enzyme (Thermo Fisher) to obtain AMPKα2(1-312)-pLenti and AMPKα2K45R-pLenti constructs. The production of lentiviral vectors (LV) batches was as previously described (Caillierez et al, 2013). Primary neurons were infected with LV at 10 DIV and were used for experiments 4 days after LV infections.

### Intracellular ATP quantification

Intracellular ATP content was quantified using the Dual-Glo^®^ Luciferase Assay System (Promega), according to the supplier’s instructions. Briefly, after treatments, primary neurons were incubated with the Dual-Glo^®^ substrate for 10 min at RT before assessing luciferase activity using Spectramax^®^ i3 (Molecular Devices).

### Phospho-Kinase array

For the phospho-kinase assay, the array membranes were incubated with cell lysates (250 μg of total proteins per array) and subsequently processed according to the manufacturer’s instructions (R&D Systems).

### Seahorse assay

For Seahorse XFe24 respirometry assay, 100,000 neuronal cells were seeded in each well. The assay medium was composed of Dulbecco’s Modified Eagle Medium base (DMEM, D-5030 Sigma) supplemented with 2 mM L-Glutamine (Invitrogen), 1.85 g/l NaCl (VWR), 3 mg/l Phenol Red (Sigma) and adjusted to pH 7.3+/-0.05. For each assay, cells were rinsed with the assay medium before being pre-incubated at 37°C without CO_2_ 20 min prior the reading. For the glycolytic test, the extracellular acidification rate (ECAR in mpH/min) was monitored during basal condition (the mixing 2:30 min, waiting 2:00 min, reading 2:30 min cycle was repeated three times) and upon the sequential addition of saturating concentration of Glucose (10 mM) (the mixing 2:30 min, waiting 2:00 min, reading 2:30 min cycle was repeated 3 times), followed by Bic/4-AP (50 μM/2.5 mM) (the mixing 1:00 min, waiting 1:00 min, reading 2:30 min cycle was repeated five times), oligomycin (Oligo 1 μM) (the mixing 2:30 min, waiting 2:00 min, reading 2:30min cycle was repeated two times), and 2DG (150 mM) (the mixing 2:30 min, waiting 2:00 min, reading 2:30 min cycle was repeated two times). Glycolysis was expressed as ECAR increase after glucose addition. The glycolytic response to Bic/4-AP was calculated as the ECAR difference before and after the injection of Bic/4-AP, glycolytic capacity was calculated as the maximum ECAR reached following oligomycin injection when compared to basal conditions before glucose injection, glycolytic reserve as the difference between ECAR increase after oligomycin addition and upon glucose injection. The spare glycolytic reserve after Bic/4-AP injection was calculated as the percentage of the glycolytic reserve spared upon response to Bic/4-AP treatment. In the mitochondrial stress test, the oxygen consumption rate (OCR in pMO_2_/min) was monitored during the basal condition (the mixing 1:30 min, waiting 2:00 min, reading 3:00 min cycle was repeated three times) and after the subsequent injection of Bic/4-AP (50 μM/2.5 mM) (the mixing 1:30 min, waiting 2:00 min, reading 2:00 min cycle was repeated three times), oligomycin (Oligo, 1 μM) (the mixing 1:20min, waiting 1:20min, reading 2:00min cycle was repeated three times), FCCP (0.5 μM) (the mixing 0:20 min, waiting 0:00 min, reading 2:00 min cycle was repeated four times), Rotenone/Antimycin A (Rot/AA 1 μM/1 μM) (the mixing 1:20 min, waiting 1:20 min, reading 2:00 min cycle was repeated three times). For mitochondrial stress test, the assay medium was supplemented with glucose (10 mM), or L-Lactate (10 mM), or NaPyr (10 mM), as specified, before the preincubation without CO_2_. Basal respiration was expressed as the difference in OCR before Bic/4-AP and after Rot/AA injection. The oxphos response to Bic/4-AP was expressed as the increase of OCR before and after Bic/4-AP addition. ATP turnover coupled to Bic/4-AP was calculated as the OCR difference before and after oligomycin injection. Maximal respiration was calculated as the OCR increase after FCCP compared to OCR upon Rot/AA injection, while spare respiratory capacity was expressed as the increase in OCR after FCCP when compared to basal respiration OCR level. Spare respiratory capacity coupled to Bic/4-AP was calculated as the percentage of mitochondrial OCR spare respiratory capacity when coupled to Bic/4-AP injection. The pH of the drug solutions were adjusted to pH 7.3+/-0.05 prior to their use.

### Metabolomics analysis

Five hundred thousand neurons were supplemented with media containing uniformly labelled ^13^C-glucose (25 mM) for 30 min. Cells were rinsed with ice-cold 0.9% NaCl solution and metabolites were extracted by adding 250 μl of a 50-30-20 solution (methanol-acetonitrile-10 mM tris HCl pH 9.4) to the cells. Plates were then incubated for 2-3 min on ice. Cells were scraped and transferred to an eppendorf tube before centrifugation at 20,000 × g for 10 min at 4°C, supernatants were then transferred to a fresh tube and stored at -80°C. Separation of metabolites prior to Mass Spectrometry (MS) measurement was performed using a Dionex UltiMate 3000 LC System (Thermo Scientific) coupled to a Q Exactive Orbitrap mass spectrometer (Thermo Scientific) operating in negative ion mode. Practically, 15 μl of the cellular extract was injected on a C18 column (Aquility UPLC^®^HSS T3 1.8μm 2.1x100mm) and the following gradient was performed by solvent A (H_2_O, 10mM Tributyl-Amine, 15mM acetic acid) and solvent B (100% Methanol). Chromatographic separation was achieved with a flowrate of 0.250ml/min and the following gradient elution profile: 0min, 0%B; 2min, 0%B; 7min, 37%B; 14min, 41%B; 26min, 100%B; 30min, 100%B; 31min, 0%B; 40min, 0%B. The column was placed at 40°C throughout the analysis. The MS operated both in full scan mode (m/z range: 70-1050) using a spray voltage of 4.9 kV, capillary temperature of 320°C, sheath gas at 50.0, auxiliary gas at 10.0. The AGC target was set at 3e6 using a resolution of 140.000, with a maximum IT fill time of 512 ms. Data collection was performed using the Xcalibur software (Thermo Scientific).

### Western blot (WB)

For WB analysis, proteins from total cell lysates were separated in 8-16% Tris-Glycine gradient gels and transferred to nitrocellulose membranes. Membranes were then blocked in 5% fat-free milk in TBS-0.01% Tween-20, and incubated with specific primary antibodies overnight at 4 °C. Proteins were thereafter detected via the use of HRP-conjugated secondary antibodies and ECL detection system (ThermoFisher Scientific).

### Statistical analysis

All statistical analyses were performed using GraphPad Prism (Prism 5.0d, GraphPad Software Inc, La Jolla, CA, USA).

## Acknowledgements

We thank the animal core facility (animal facilities of Université de Lille-Inserm) of “Plateformes en Biologie Santé de Lille” as well as C. Degraeve, M. Besegher-Dumoulin, J. Devassine, R. Dehaynin and D. Taillieu for animal care. The authors thanks the behavioral exploration platform for rodent (Federation of Neurosciences, Univ. Lille, France). pAMPK alpha2 WT and pAMPK alpha2 K45R were a gift from Morris Birnbaum (Addgene plasmids# 15991 and 15992).

This work was supported by the French Fondation pour la cooperation Scientifique – Plan Alzheimer 2008-2012 (Senior Innovative Grant 2013) to VV, by the Fondation Vaincre Alzheimer (n°FR-16071p to VV), and in part through the Labex DISTALZ (Development of Innovative Strategies for a Transdisciplinary Approach to Alzheimer’s disease). TA and DB are supported by the Fonds voor Wetenschappelijk Onderzoek (FWO) Flanderen, project G0D7614N. MD holds a doctoral scholarship from Lille 2 University.

---

**Figure EV1.**
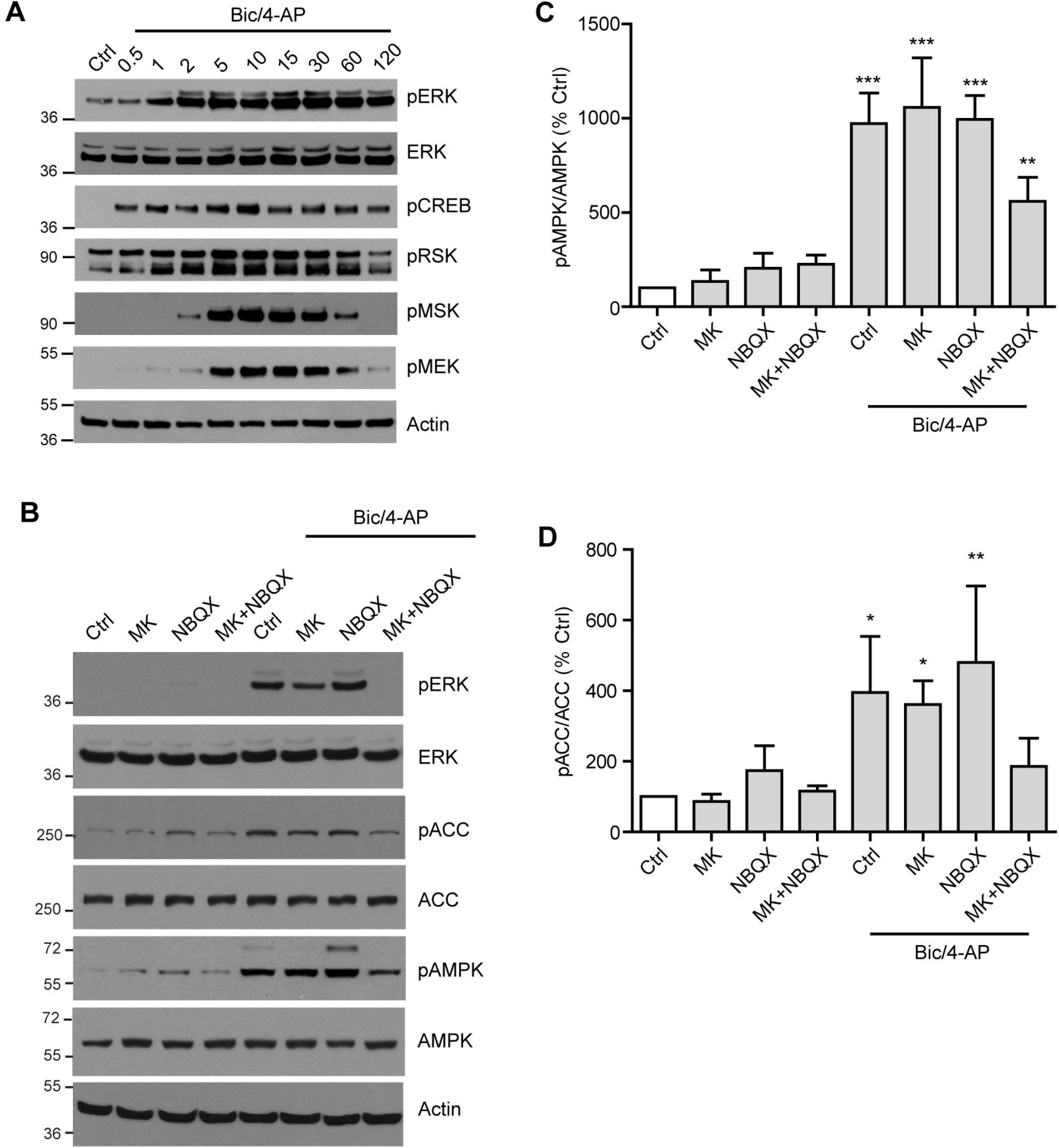
AMPK activation following SA is dependent on NMDA and AMPA receptors activation. A Primary neurons were treated with Bic/4-AP for the indicated times and the activation of the MAPK pathway was monitored by immunoblotting with anti-phosphorylated ERK, CREB, RSK, MSK, MEK and total ERK and Actin antibodies. Results are representative of at least 3 experiments. B Primary neurons were pre-treated for 20 min with the NMDA receptors and AMPA receptors antagonists MK-801 (25 μM) and NBQX (20 μM), respectively. After pre-treatment, neurons were stimulated with Bic/4-AP for 10 min. Activation of ERK and AMPK were monitored by immunoblotting with anti-phosphorylated ERK and total ERK and anti-phosphorylated AMPK and ACC, total AMPK, ACC, and Actin antibodies. Results are representative of at least 5 experiments. C-D Quantifications of WB as shown in B displaying the ratios phosphorylated AMPK/total AMPK (pAMPK/AMPK) and phosphorylated ACC/total ACC (pACC/ACC) expressed as percentage of control (n=3-5). Results show mean ± SD. One-way ANOVA followed by Bonferroni’s post-test was used for evaluation of statistical significance. * p < 0.05, ** p < 0.01, *** p < 0.001).

**Figure EV2.**
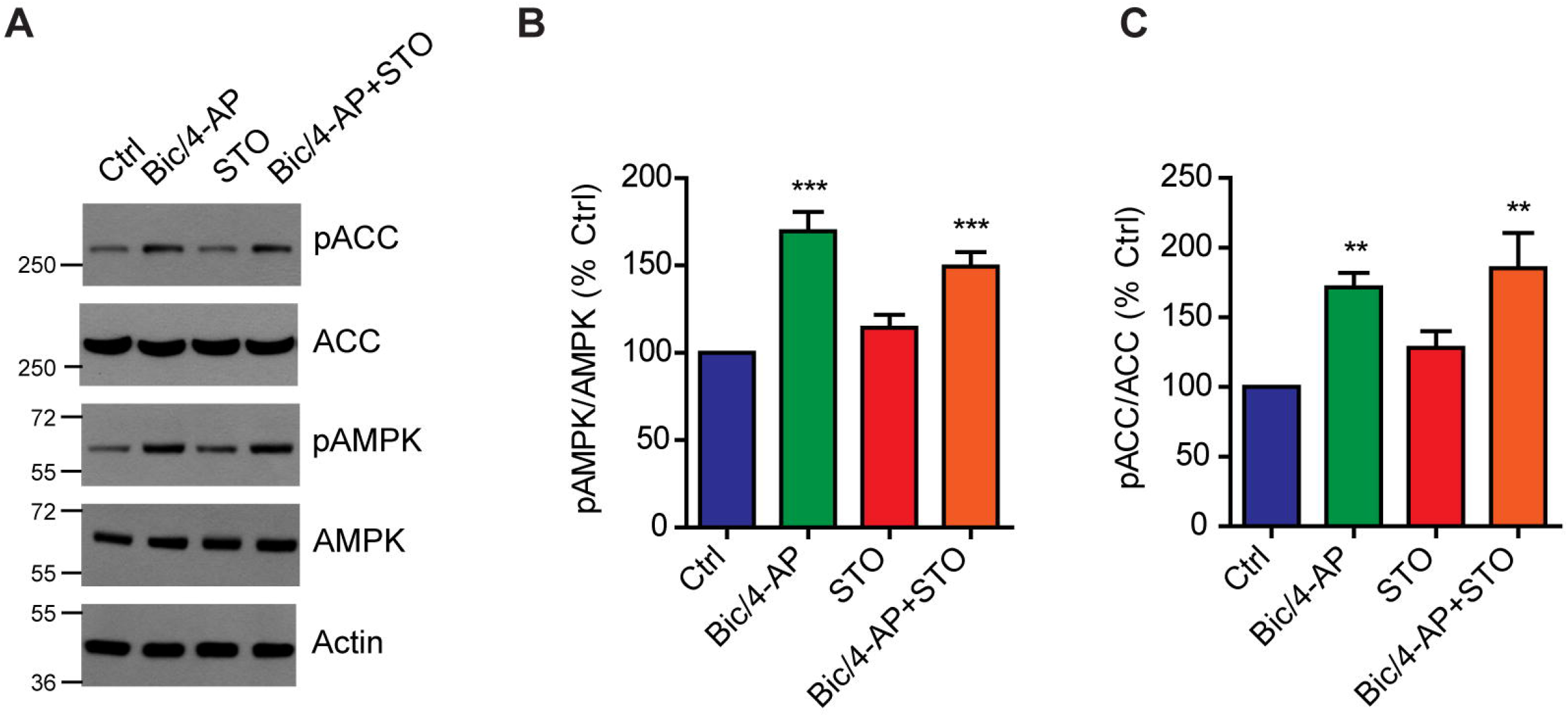
SA induced activation of AMPK is not mediated by CaMKKβ. A Primary neurons treated with Bic/4-AP (10 min) after 20 min pre-treatment or not with the CaMKKβ specific inhibitor STO-609 (STO, 10 μM) were subjected to immunoblotting with anti-phosphorylated AMPK and ACC, total AMPK and ACC, and actin. Results are representative of at least 5 experiments. B-C Quantifications of WB as in A showing the ratios phosphorylated AMPK/total AMPK (pAMPK/AMPK) and phosphorylated ACC/total ACC (pACC/ACC) expressed as percentage of control (n=5). Results show mean ± SD. One-way ANOVA followed by Bonferroni’s post-test was used for evaluation of statistical significance. * p < 0.05, ** p < 0.01, *** p < 0.001).

**Figure EV3.**
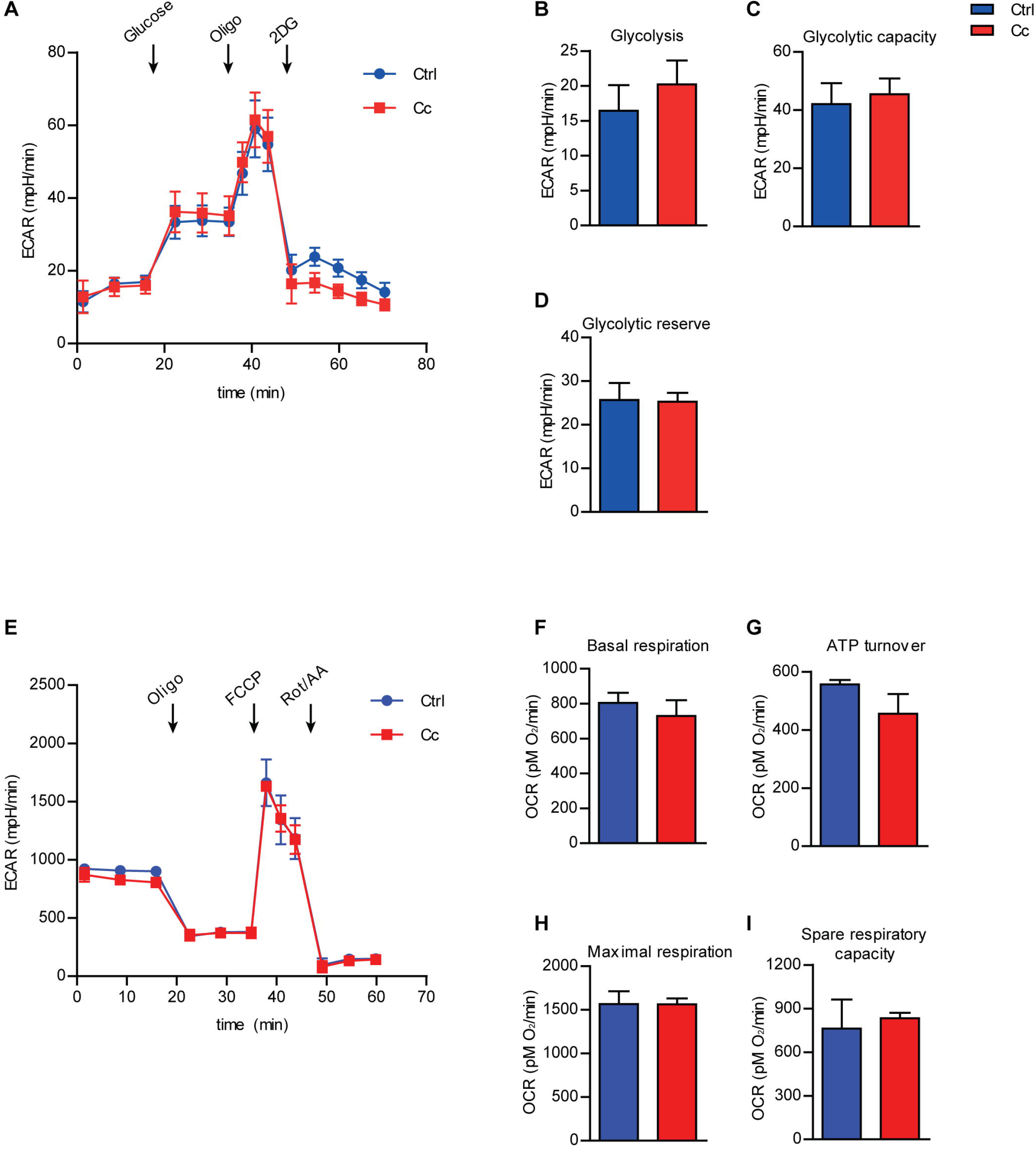
Effect of Compound C on glycolytic and mitochondrial oxidative phosphorylation profiles in primary neurons. A-I (A) Extracellular acidification rate (ECAR) and (E) oxygen consumption rate (OCR) measured using the Seahorse technology in the presence or absence of the AMPK inhibitor Compound C (Cc, 10 μM). (A) ECAR profile is monitored under basal conditions and upon the sequential injection of saturating concentration of glucose, Oligo and 2DG as indicated by arrows. (E) OCR profile monitored under basal condition and following the sequential injection of Oligo, FCCP, and Rot/AA as indicated by the arrows. ECAR and OCR are indicators of glycolysis and mitochondrial respiration, respectively. (B) Basal glycolysis, (C) glycolytic capacity, and (D) spare glycolytic reserve expressed as mpH/min. (F) Basal respiration, (G) ATP turnover stimulation, (H) maximal respiration, and (I) spare respiratory capacity expressed as pMO_2_/min. Results are representative of at least 3 independent experiments. Results show mean ± SD. One-way ANOVA (B-D, F-I) followed by Bonferroni’s post-test were used for evaluation of statistical significance. * p < 0.05, ** p < 0.01, *** p < 0.001.

**Figure EV4.**
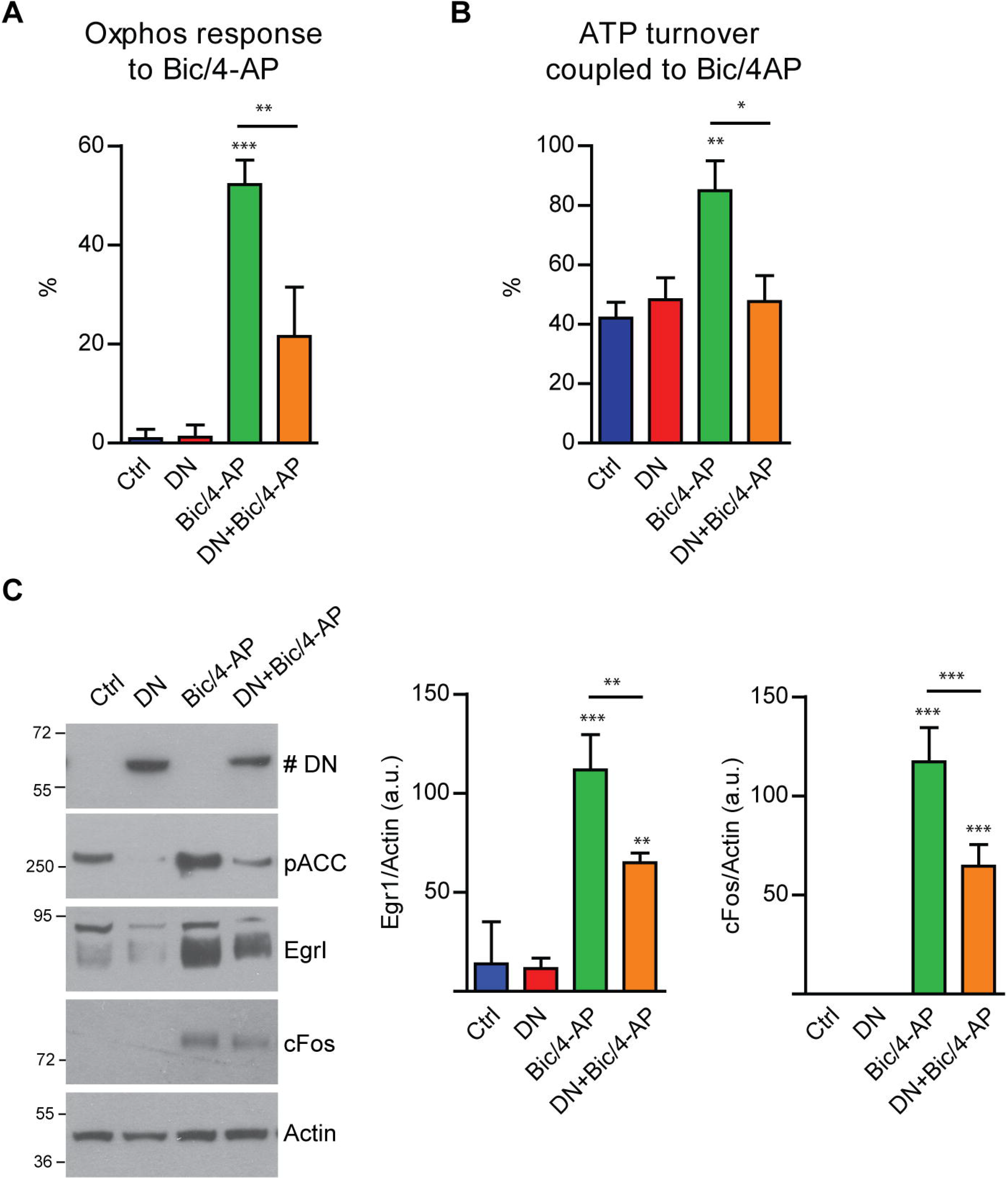
AMPK inhibition using a dominant negative construct repress the metabolic response to SA and the expression of IEGs. A-B OCR response to Bic/4-AP stimulation (A) and ATP turnover coupled to Bic/4-AP (B) were evaluated using the Seahorse technology in primary neurons expressing a dominant negative (DN) form of AMPK. Results are expressed as percentage of the first respiration point (n=3). C Immunoblotting with anti-myc (# DN labels the overexpressed DN-AMPK construct), Egrl, cFos, and Actin in lysates obtained from 15 DIV neurons stimulated with Bic/4-AP for 2 hr, 4 days after infection with DN-AMPK. Results are representative of 3 experiments. Quantification of WB showing the expression of the IEGs Egrl and cFos (n=3). Results show mean ± SD. One-way ANOVA followed by Bonferroni’s post-test was used for evaluation of statistical significance. * p < 0.05, ** p < 0.01, *** p < 0.001.

**Figure EV5.**
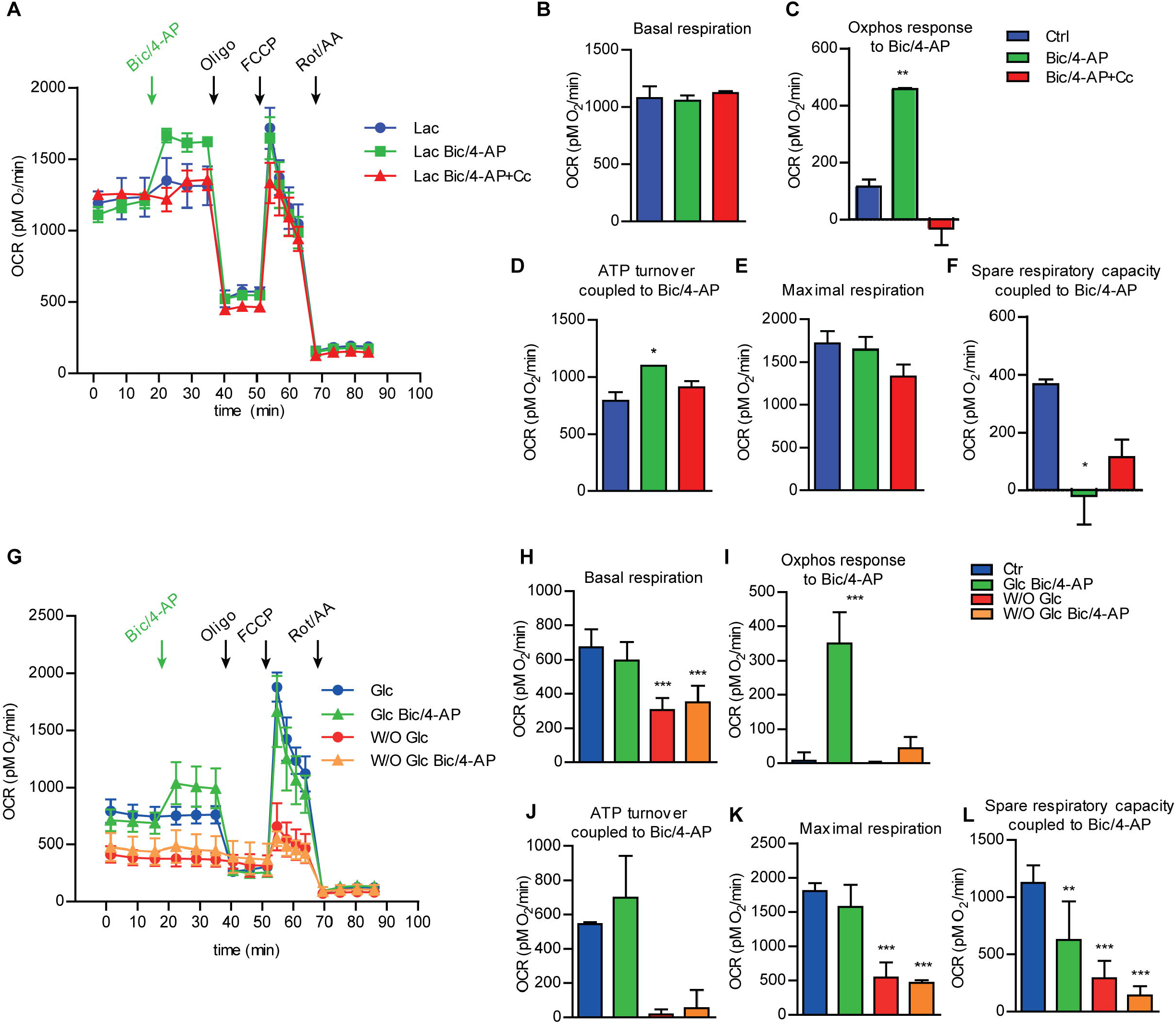
Utilization of mitochondrial alternative substrates by primary neurons in response to SA. A Oxygen consumption rate (OCR) measured using the Seahorse technology after Bic/4-AP stimulation (green arrows) in the presence or absence of L-Lactate (Lac, 10 mM). OCR profile was monitored under basal conditions and following the sequential injection of assay medium, in Ctrl group, or Bic/4-AP, in the stimulated groups, Oligo, FCCP, and Rot/AA as indicated by the arrows. B-F (B) Basal respiration, (C) oxphos response to Bic/4-AP, (D) ATP turnover coupled to Bic/4-AP stimulation, (E) maximal respiration, and (F) spare respiratory capacity coupled to Bic/4-AP expressed as pMO_2_/min. G-L (G) Oxygen consumption rate (OCR) measured using the Seahorse technology after Bic/4-AP stimulation (green arrows) in the presence (Glc, 10 mM) or absence (W/O Glc) of glucose. OCR profile was monitored under basal conditions and following the sequential injection of assay medium, in Ctrl group, or Bic/4-AP, in the stimulated groups, Oligo, FCCP, Rot/AA as indicated by the arrows. (H) Basal respiration, (I) oxphos response to Bic/4-AP, (J) ATP turnover coupled to Bic/4-AP stimulation, (K) maximal respiration, and (L) spare respiratory capacity coupled to Bic/4-AP expressed as pMO_2_/min. Results are representative of at least 3 independent experiments and show mean ± SD. One-way ANOVA followed by Bonferroni’s post-test was used for evaluation of statistical significance. * p < 0.05, ** p < 0.01, *** p < 0.001.

